# Structural insights into the inhibition of sickle hemoglobin polymerization by asymmetric hemoglobin tetramer HbFS (α_2_γβ_S_)

**DOI:** 10.64898/2026.06.03.729826

**Authors:** Aastha baliyan, Neha Yadav, Nihar Ranjan Mishra, Santosh Kumar Mondal, Kalyan Goswami, Jayeeta Bhowmick, Amit Kumar Mandal

## Abstract

Sickle cell disease (SCD) is caused by a single amino acid substitution in the β_S_ globin chain at 6th position (6E→V). This results in polymerization of deoxy state of sickle hemoglobin (HbS), followed by its precipitation and subsequent sickling of erythrocytes. These deformed cells can block small capillaries (vaso-occlusion), causing cardiovascular complications, ultimately leading to ischemia-reperfusion injury, severe oxygen deficiency, and progressive systemic damage. Occasionally, patients with SCD have been observed to produce exorbitantly high levels of fetal hemoglobin (HbF), which has been linked with the inhibition of HbS polymerization. One of the effects of hydroxyurea, the most commonly used therapeutic for SCD, is to elevate HbF levels. However, the mechanism of inhibitory role of HbF on HbS polymerization is largely unknown. This study attempts to gain insights into the mechanisms involved in this process by means of native mass spectrometry, ion mobility mass spectrometry, and hydrogen deuterium exchange-based mass spectrometry (H/DX-MS). The conformational flexibility of asymmetric hemoglobin, HbFS (α_2_γβ_S_), for the observed regions in the tetrameric molecule appears to be more in the deoxy state as compared to the oxy state, eventually leading to reduced polymerization of sickle hemoglobin in patients with SCD that express elevated HbF levels.

## Introduction

Sickle cell disease (SCD) is a genetic disorder caused by an amino acid substitution at the 6th position of the β globin chain (6E→V) (1). This variant human hemoglobin, known as HbS, promotes intermolecular hydrophobic interactions, leading to the formation of long polymer chains in the deoxy state (2–4). The isopropyl group β_S_Val6 residue of sickle hemoglobin tetramer, in deoxy state, forms a hydrophobic patch which binds to the hydrophobic pocket of another tetramer, whose hydrophobic patch binds the hydrophobic groove of a third tetramer, and so on (2). The polymerization of HbS in deoxy state is entropically driven and takes place in two phases: a delay period followed a rapid polymerization process (5). During the delay time, homogeneous polymerization of about 15–30 HbS tetramers results in the formation of a single nucleus (5–7). These nuclei then undergo heterogeneous polymerization, leading to formation of a double-stranded polymer that elongates as the polymerization proceeds and the fiber growth continues approximately up to 250 tetramers (8). Seven such double-stranded polymers associate to form a fourteen-stranded insoluble fibrous polymer (2). Precipitation of these fibers (n>14) is sufficient to deform the shape of Red Blood Cells (RBCs) into the characteristic sickle shape observed in SCD, thus giving the disease its name (9–11). These deformed cells ultimately lead to several health complications resulting from membrane damage, hemolysis, vaso-occlusion, ischemia-reperfusion injury, inflammation, cardiovascular complications, and progressive multiorgan dysfunction (12–14).

There are several factors that can modulate the polymerization of HbS (15,16). One of the best-known factors that can reduce the polymerization of HbS is the presence of another hemoglobin variant: fetal hemoglobin (HbF) (17–21). After birth, the level of HbF reduces and is eventually replaced by adult hemoglobin (HbA), and is therefore present only in trace amounts in adults (<1%) (22). However, patients with SCD and/or β-thalassemia often express relatively high levels of HbF (17,21,23). Occasionally, it has been observed that a very few patients with SCD produce exorbitantly high levels of fetal hemoglobin (HbF). This abnormally high expression of HbF in adults is clinically very important because it reduces HbS polymerization, thereby prolonging survival (3,17). Epidemiological studies have consistently shown that higher HbF levels frequently correlate with nearly asymptomatic disease (17,21,24). Hydroxyurea, a common therapeutic used to treat SCD, acts, in part, by elevating HbF levels (25–29).

Despite decades of study, the molecular insights into the inhibitory nature of HbF on HbS polymerization are still unknown. Molecular crowding may be one reason HbF effectively reduces the likelihood of HbS tetramer nucleation, thereby delaying polymerization (3,17,19,30,31). Although HbF by itself doesn’t participate in the polymerization, reports have described the presence of asymmetric tetrameric hemoglobin (HbFS) molecules in patients with elevated HbF levels (19,30). HbFS consists of two α globin chains and one β globin chain from each of the sickle and gamma variants. However, the crystal structure of the HbFS (α_2_β_S_γ) tetramer has not yet been resolved. Thus, while the elevation of HbF remains the most validated therapeutic strategy for SCD, the precise structural and conformational dynamics underlying the mechanism by which HbF modulates HbS polymerization are largely unknown to date (29).

In this study, we analyzed a sample from a 2-year-old child presenting with homozygous sickle and β thalassemia trait. HbF levels in this child were 40.1% of total hemoglobin, as reported by cation-exchange HPLC-based hemoglobin analysis in the diagnostic reports. We studied conformational changes in asymmetric hemoglobin tetramers to explore the underlying mechanism of sickle hemoglobin polymerization inhibition, using a mass spectrometry-based platform.

## Materials and methods

### Materials

Highly pure crystalline pepsin, Deuterium oxide (99.9%), α-cyano-4-hydrocinnamic acid (CHCA), LC-MS grade water, ammonium acetate (>99%), ammonium bicarbonate, LC-MS grade trifluoroacetic acid (TFA), sodium chloride (>99%), Red phosphorus.

### Sample collection

We received 1.2 ml of the leftover blood samples from a 2-year-old child and from parents of the child who visited AIIMS Kalyani for a general health checkup. The sample was collected at AIIMS Kalyani hospital for further analysis, with the patient’s prior consent and the study was approved by the IEC of both AIIMS Kalyani and IISER Kolkata. The patient’s RBCs were observed to be teardrop-shaped and elongated. HPLC analysis of the patient’s hemoglobin profile at the diagnostic facilities at AIIMS Kalyani revealed the presence of HbS, HbF, and HbA at 51.1, 40.1, and 8.8%, respectively. The patient’s mother (HbA-58.6%, HbA_2_-2.9%, HbS-38.1%) was diagnosed as a carrier of SCD, and the father (HbA-83.6%, HbA2-5.7%, HbF-10.5%) was diagnosed with thalassemia trait, as determined by cation-exchange HPLC analysis.

### Ex vivo RBC sickling

The sickling experiment was carried out using the method reported by Young-Zoon Yoon et al., with slight modifications (32). In brief, suspensions of RBCs in 0.9% saline were deoxygenated using 2 mg/ml sodium hydrosulfite. A drop of the suspension was placed on the slide, and the coverslip was placed over it and sealed with nail polish to prevent air exposure. The slide was subjected to morphological analysis using a phase contrast microscope. Images were captured using a camera attached to the microscope with TCapture software. Several images were captured, and the percentage of sickle RBCs was determined using the following equation:

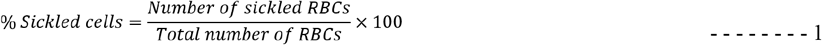

### Sample processing

Whole blood samples were centrifuged at 3000 rpm for 5 min at room temperature. Following centrifugation, the plasma and buffy coat layers were carefully removed, and the red blood cell (RBC) pellet was retained. The RBC pellet was washed three times with three volumes of normal saline (v/v), with each wash followed by centrifugation at 3000 rpm for 5 min at room temperature. The washed RBC pellet was lysed by adding eight volumes of ice-cold water, followed by a brief vortex. The lysate was subsequently centrifuged at 12,000 rpm for 15 min at 4 °C to separate cellular debris from the hemolysate. The resulting supernatant was collected, and the hemoglobin concentration was determined spectrophotometrically at 548 nm using a UV-Visible spectrophotometer (Shimadzu). The prepared hemolysate was aliquoted and snap frozen in liquid nitrogen. The samples were stored at -80 °C until use.

### Native ESI-MS-based mass spectrometric analysis

The hemolysate was dialyzed overnight against a 10 mM ammonium acetate buffer (pH 7.4) and diluted to a final concentration of 10 µM. Samples were introduced into a Waters Synapt G2-Si mass spectrometer (Waters, UK) using a micro-ESI Z-spray source in direct infusion mode at a flow rate of 2 µL/min. Prior to analysis, the mass spectrometer was calibrated using 2 mg/mL sodium iodide in the m/z range 50-5000. The source temperature was maintained at 37 °C to retain the non-covalent interaction within the native tetrameric structure of hemoglobin. Instrument parameters were set as follows: capillary voltage 2 kV, trap CE (Collision Energy) 15 V; transfer CE 2 V; desolvation temperature 250 °C; and desolvation gas flow 500 L/h. Data were acquired in positive-ion mode over an m/z range of 700–4000 at an acquisition rate of 1 spectrum/s, with analyzer in V-mode. The quad profile was set to manual as follows: 1600 (M1), 3 (D1), 0 (R1), 400 (M2), 37 (D2), 60 (R2), and 4000 (M3).

The data analysis was performed using Masslynx 4.1 for baseline correction and smoothening of the data. For deconvolution of the spectrum in order to obtain the neutral masses of the molecules, MaxEntI algorithm was used.

### Ion mobility based mass spectrometric analysis

The hemolysate samples were analyzed in ion mobility mode by direct infusion using a micro-ESI Z-spray source coupled to a Waters Synapt G2-Si mass spectrometer under the same instrumental conditions described for native mass spectrometry in the previous section. Prior to analysis, the mass spectrometer was calibrated using 2 mg/mL sodium iodide in the m/z range 50-5000. For ion mobility separation, the helium gas flow was maintained at 45 mL/min, while the IMS gas flow was set to 30 mL/min. The ion mobility parameters were configured as follows: trap wave height 4 V; trap wave velocity 311 m/s; IMS wave velocity 950 m/s; IMS wave height 30 V; transfer wave velocity 382 m/s; and transfer wave height 40 V.

The acquired data were processed using DriftScope v2.8 (Waters, UK). Peak detection was performed in non-chromatographic mode at a resolution setting of 10,000 and a minimum intensity threshold of 1 × 10^4^ counts. The ion mobility data were calibrated using previously reported reference collision cross-section (CCS) values for HbS conformational populations (33). Charge states corresponding to the different hemoglobin populations were assigned to calculate the respective CCS values. The resulting CCS values obtained from individual tetramer charge states were averaged, followed by averaging across duplicate experimental sets prior to further analysis.

### Hydrogen Deuterium Exchange-based Mass Spectrometry (H/DX-MS)

300 µM hemolysate was diluted 15-fold with 150 mM ammonium bicarbonate/D_2_O buffer (pD 7.8), and exchange kinetics were performed at 25 °C for 3 hours. At different time intervals, a 10 µL aliquot was removed and added to 90 µL of ice-cold aqueous 0.1% TFA (pH 2.5) to quench the H/DX exchange kinetics. Aliquots of quenched isotope-exchanged hemoglobin molecules were digested *in situ* by treating with pepsin (1:10 mol/mol). The digestion was carried out for 5 min at 0 °C, pH 2.5. A 1:1 mixture of CHCA (α-Cyano-4-hydroxycinnamic acid) matrix and solution was made, and 0.5 µL of it was spotted on the MALDI plate. The spot was immediately dried in a desiccator to minimize back-exchange, and profile data were acquired in positive-ion, sensitivity mode over the mass range 700-4000 m/z using a Synapt G2-Si (Waters, UK) system with a MALDI source. The system was calibrated with red phosphorus prior to data analysis.

The experimental data were pre-processed by baseline correction and smoothening using the Savitzky-Golay algorithm using Masslynx™ software. The data for individual peptides was exported to an Excel sheet and analyzed using HX-Express v3 (34). For detecting the peptide envelope, the charge state was set to 1+, and the distribution width was set to 20% of the Base Peak Intensity (BPI). For converting the profile data to centroid data, the peak tolerances were kept at 1% of BP to analyze the data to individual peaks, consisting of m/z values and corresponding intensities of different time points.

In H/DX-MS, the uptake of deuterium with increasing time results in a right shift in the isotopically distributed peaks, thereby increasing the isotope-averaged centroid mass of the peptide isotopic peak envelope. These isotope-averaged centroid masses were used to calculate the relative deuterium uptake “D(t)”, which was then plotted against time (35).

The observed deuterium uptake was calculated from the isotope-averaged centroid mass of the signal envelope at different time points with the equation:

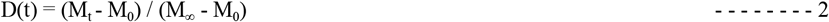

In this equation, M_t_ is the observed isotope average centroid mass of the peptide molecular ion at time ‘t’; M_0_ is the isotope-averaged centroid mass of the peptide molecular ion at time (t=0) and M_∞_ is isotope-averaged centroid mass of the peptide molecular ion at time (t=∞).

For any experimental molecule, the observed deuterium uptake can be used to measure the kinetics according to the method of initial rates, using the following equation:

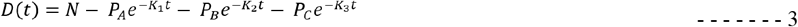

In this equation, N denotes the total exchangeable amide hydrogens in the peptide. P_A_, P_B_ and P_C_ denote the hydrogens with a fast exchange rate, intermediate exchange rate and hydrogens with a slow exchange rate respectively. k_1_, k_2_ and k_3_ are the average rate constants for the P_A_, P_B_, and P_C_.

Using the best fit analysis in MS Excel solver to fit the observed values from equation 1, by varying the populations and rate constants of equation 2, such that the sum of squared residues (SSR) is minimised, we obtained the values for P_A_, P_B_, P_C_, k_1_, k_2_, and k_3_. The SSR was calculated by the following equation:

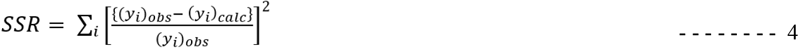

where (y_i_)_obs_ and y_i_)_calc_ were calculated from equation 1 and equation 2, respectively. the best-fit was subjected to following constraints:

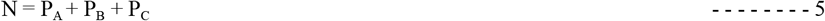

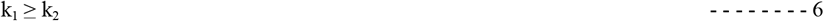

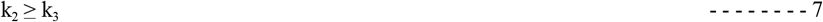

Using these six values (P_A_, P_B_, P_C_, k_1_, k_2_, and k_3_), it is possible to investigate the conformational changes associated with the transition of the deoxy to the oxy state of the HbFS sample. The algebraic sum of the differential exchange rates across three groups of amide hydrogens between two states of the experimental protein mirrors the change in the molecule’s conformational dynamics associated with the biological event. A positive sign indicates increased flexibility and a negative sign indicates increased rigidity in the conformational dynamics associated with the state change of the protein molecule (35).

## Results and discussion

The sample from the 2-year-old child co-expressing homozygous sickle (51.1% HbS) and thalassemia trait showed abnormally high expression of the HbF tetramer (40.1%) as quantified by clinical analysis (Supplementary Figure 1). The microscopy assay showed very little sickling under deoxy conditions monitored for 2 hours (Figure 1(a-b)). The proportion of sickled cells following deoxygenation was calculated by excluding cells that were already sickled prior to deoxygenation. At 0 hr, 1.75 ± 0.045% of RBCs were observed to be already sickled due to the irrevocable sickling of certain RBCs (Figure 1a). In addition to sickle cells at 0 hr, under the deoxy conditions only 0.945 ± 0.19% of cells underwent sickling after 2 hours (Figure 1b). Figure 1c depicts the fraction of sickled cells at 0 hr and 2 hr in the deoxy state, calculated according to equation 1. This very low percentage of sickling might be attributed to the presence of exorbitantly high levels of HbF in the sample, which is known to have an anti-sickling effect by inhibiting HbS polymerization (17,18,20,21). Cells that undergo sickling may be due to a differential distribution of HbF, since some cells may have a low HbF concentration (36).

**Figure 1:**
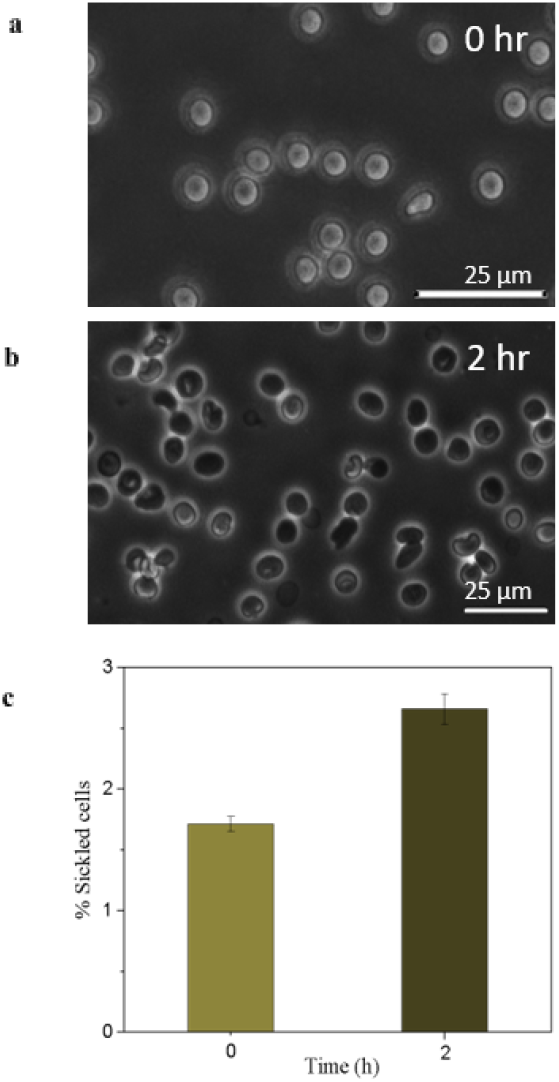
Morphology of the RBCs at 0 h (a) and 2 h (b) time points after inducing deoxy conditions was recorded at 40X magnification; (c) The percentage of sickled-shaped RBCs at 0 and 2 h time points after inducing deoxy conditions.

The hemolysate, obtained from the sample, was analyzed using native mass spectrometry. Figure 2a shows the respective native mass spectrum which shows the presence of signals corresponding to monomers of globin chains, dimers formed by the association of globin monomers, and different tetrameric species that are present in the human hemoglobin sample under investigation. Figure 2b shows the signals corresponding to different states of α, β_S_, and γ (γ_G_ and γ_A_) globin chain monomers. Inset shows the deconvoluted masses of all monomeric globin chains. However, the presence of the normal β globin chain was not observed. Figure 2c shows that the α chain forms dimers with β_S_ and γ chain, resulting in the formation of different tetrameric hemoglobins, including the asymmetric hemoglobin HbFS. Figure 2d illustrates the signals corresponding to three charge states (17+, 18+, and 19+) of different tetrameric hemoglobin populations. Upon deconvolution (Inset, figure 2c), these signals reveal three major tetrameric species: HbS (α_2_(β_S_)_2_), HbF (α_2_γ_G_γ_A_), and HbFS (α_2_γ_G_β_S_). Although we could not annotate the tetrameric species α_2_γ_A_β_S_, this does not exclude the possibility of γ_A_ to participate in the formation of asymmetric hemoglobin. Since the molecular mass difference between both the γ globin chains is only 13 Daltons, the α_2_γ_G_β_S_ and α_2_γ_A_β_S_ tetramers may not have been resolved in the present setup. For this reason, we are hereby denoting the asymmetric hemoglobin as HbFS (α_2_γβ_S_).

**Figure 2:**
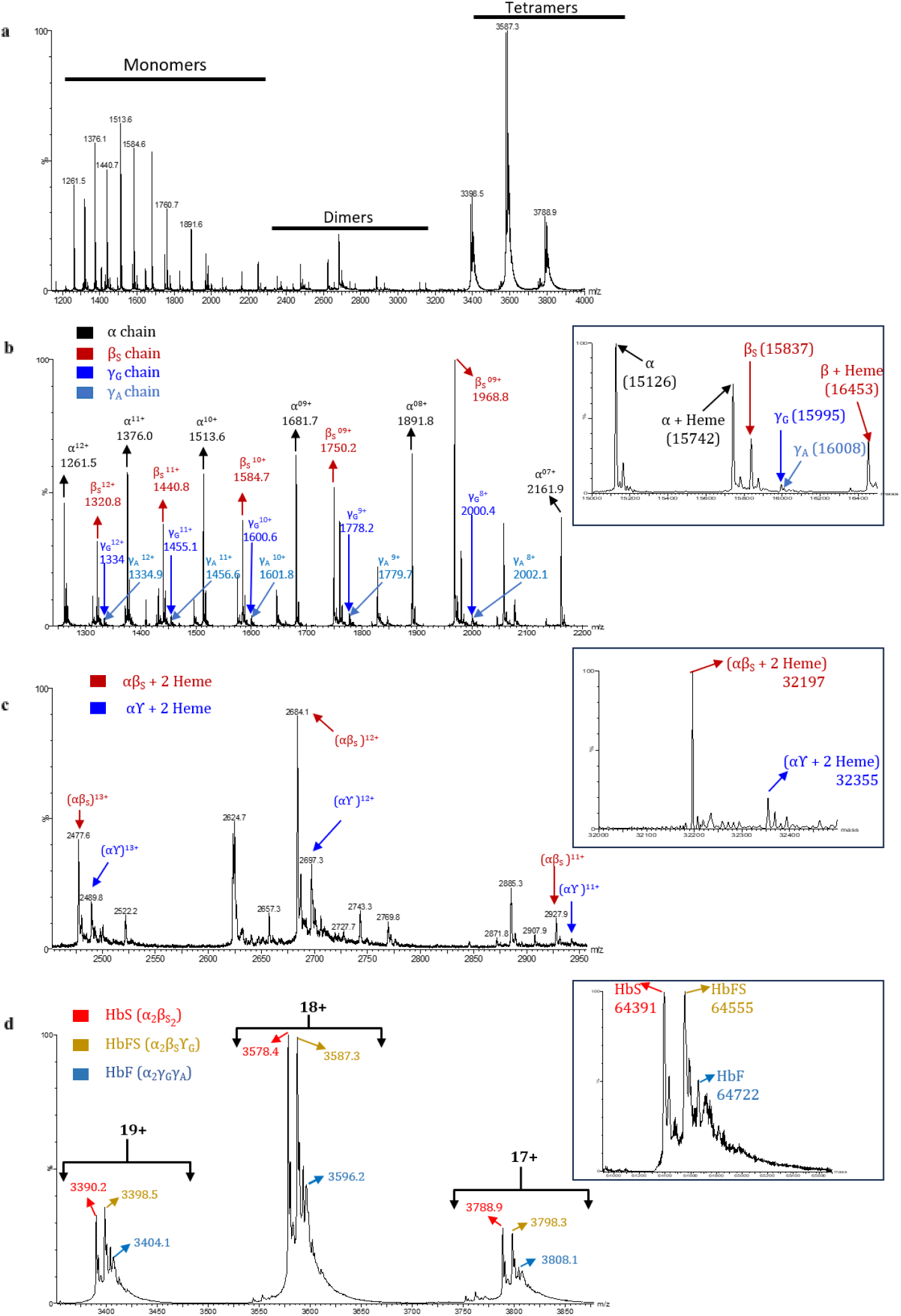
10 µM hemolysate was dialyzed against 10 mM ammonium acetate buffer (pH 7.4) and analyzed using native ESI-MS on the micro Z-spray ESI source connected to the Synapt G2-Si mass spectrometer. (a) The native mass spectrum in the mass range of 500-4000 m/z shows the presence of monomers, dimers, and tetramers of human hemoglobin in the sample. (b) The monomers were observed in the mass range of 1200-2200 m/z. The inset shows the deconvoluted mass spectrum of the monomers, indicating the presence of different globin chains: α, β_S_, and γ (γ_G_ and γ_A_). (c) Three charge states (11+, 12+ and 13+) of the dimers αβ_S_ and αγ associated with two heme moieties each were observed, inset shows their respective deconvoluted masses (d) Three charge states (17+, 18+, and 19+) of the tetramers were observed in the mass range of 3300-3900 m/z. The inset shows the presence of different tetramers: HbS (α_2_β_S(2)_, 64391 Da), HbFS (α_2_γ,_G_β_S_, 64555 Da), and HbF (α_2_γ_G_γ_A_, 64722 Da).

The elevated levels of γ globin chain monomer likely compete with the β_S_ globin chain in the association with α globin chain during the formation of the tetrameric hemoglobin molecule. High abundance of HbFS tetramers in the experimental sample indicates the comparable compatibility of the β_S_ and γ (γ_G_ and γ_A_) globin chains with α globin chain. From a structural perspective, the presence of γ globin chains reduces the likelihood of HbFS tetramers to participate in the polymerization. This might be due to the unavailability of the hydrophobic patch in the γ globin chain, which subsequently reduces the propensity of HbFS tetramers to participate in the elongation of polymer strands. Consequently, the presence of γ chains is likely to disrupt the intermolecular and intramolecular interactions that are required to stabilize the polymer fibrils. It supports the proposed role of fetal hemoglobin in modulating HbS polymerization through molecular crowding and dilution effects (31).

Despite such pronounced effects of HbFS tetramer on the polymer formation, the measured cross-sectional area of HbS, HbFS, and HbF tetramers does not vary significantly from each other, as evidenced by the similar cross-sectional area of the constituting dimers,(αβ_S_ and αγ_G_) (Figure 3a-d). This indicates that the overall dimensions of the dimers are largely conserved in order to be incorporated into the tetramer. This finding suggests that the inhibitory effects of HbFS on HbS polymerization are independent of the global rearrangements or steric alterations in the overall structure as compared to the HbS tetramer. It might indicate that the inhibitory effects of HbFS on HbS polymerization may be attributed to subtle differences in the surface chemistry, intermolecular interactions, and/or local conformational dynamics rather than global differences in the tetrameric structure. These localized changes may modulate the lock-and-key mechanism that mediates the association of HbS tetramers.

**Figure 3:**
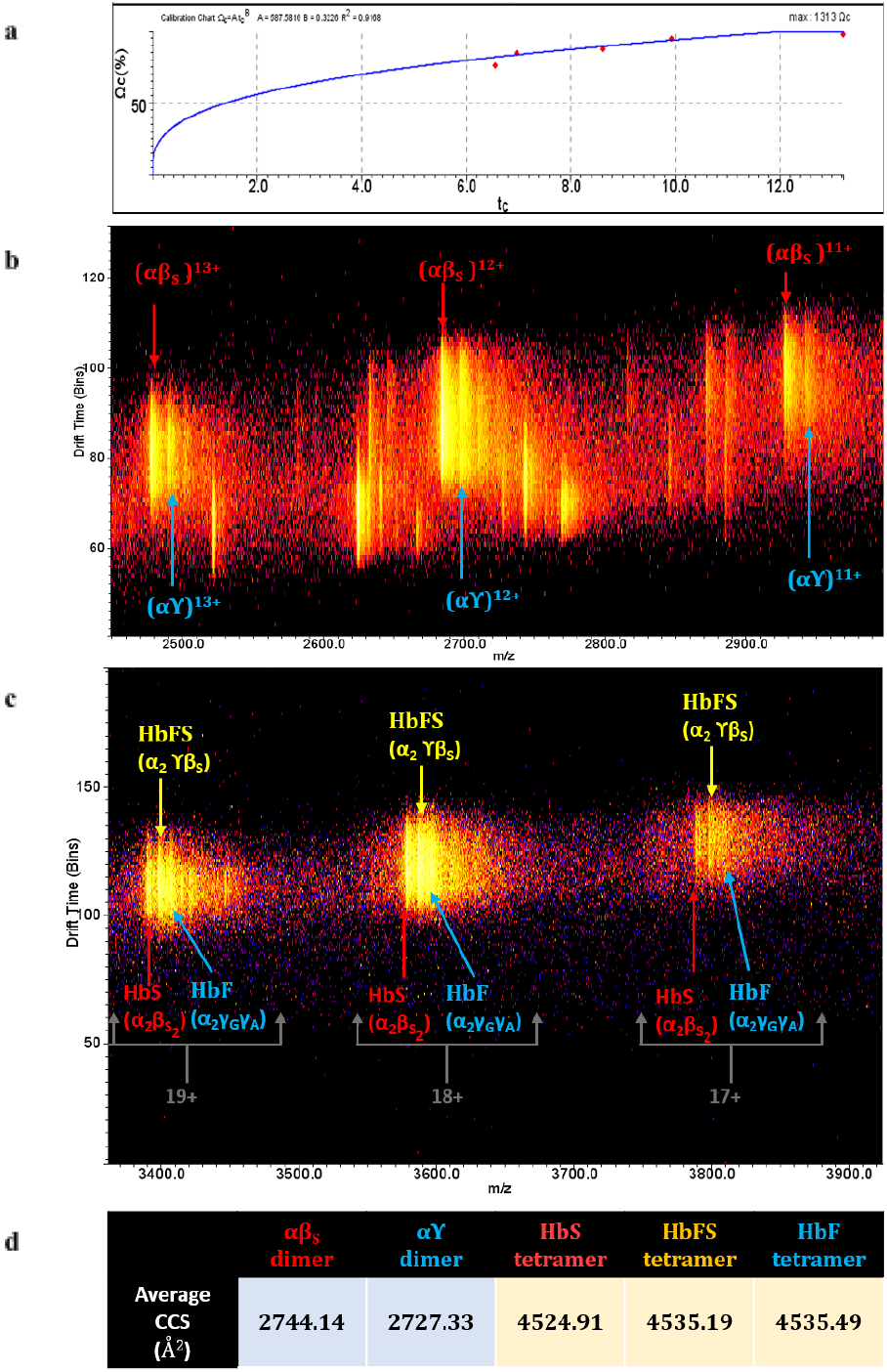
Ion mobility-based separation of the tetrameric populations: (a) The calibration curve of the recorded data against the reference values of HbS tetramers gave an R^2^ value of 0.9168. (b) The ion mobility spectrum shows that for every charge state, dimers αβ_S_ and αγ have similar drift times. (c) The ion mobility spectrum shows different tetramers (HbS, HbFS, and HbF) in three different charge states (17+, 18+, and 19+) with similar drift time. (c) The average CCS values of three different dimers (αβ_S_ and αγ) and tetramers (HbS, HbFS, and HbF).

H/DX-MS was employed in order to investigate changes in confirmation of tetrameric HbFS associated with the transition between the oxy and deoxy states. The hemolysate in two states (deoxy and oxy) was exposed to deuterium oxide (D_2_O) in separate sets, and the isotope exchange (H/DX) kinetics were monitored for 180 minutes and analyzed using MALDI mass spectrometry platform. Depending on solvent accessibility and conformational flexibility, the isotope-averaged centroid mass of a proteolytic peptide increases with increasing exposure time to D_2_O. Figure 4 (a-g) shows the rightward shift in the isotopic peak envelopes of two representative peptides from each of the α and β_S_ globin chains in oxy and deoxy states. We measured the differences in the relative deuterium uptake of eight different peptides comprising four from the α chain and four from the β_S_ chain (Supplementary Figure 2) in their oxy and deoxy states (Figure 4 h-p). The predicted deuterium uptake (equation 3) was fitted to the observed deuterium uptake (equation 2) by using best-fit analysis to reduce the SSR to 0 by varying the populations and rate constants, for each peptide in both conformational states (Figure 5 a-h). The analysis provided us with the kinetic parameters (P_A_, P_B_, P_C_, k_1_, k_2_, and k_3_) for the deuteration kinetics in each peptide in both conformational states. The kinetic parameters obtained were then used to calculate conformational flexibility in two states (Supplementary Figure 3). Table 1 shows the difference in the summation of rates of isotope exchange (k_1_, k_2_, k_3_) for three different groups (P_A_, P_B_, P_c_) of backbone amide hydrogens in both oxy and deoxy states.

**Figure 4:**
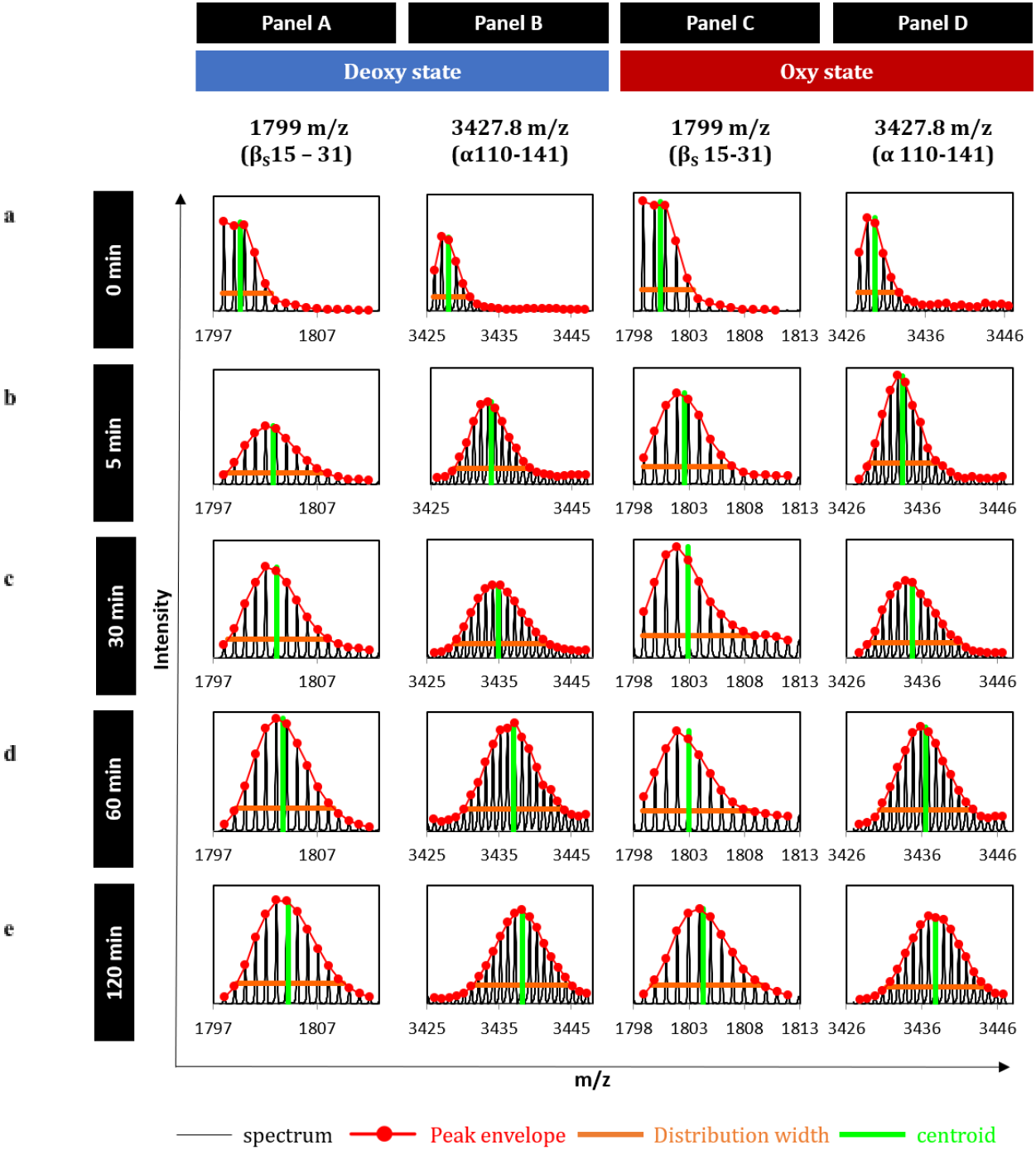
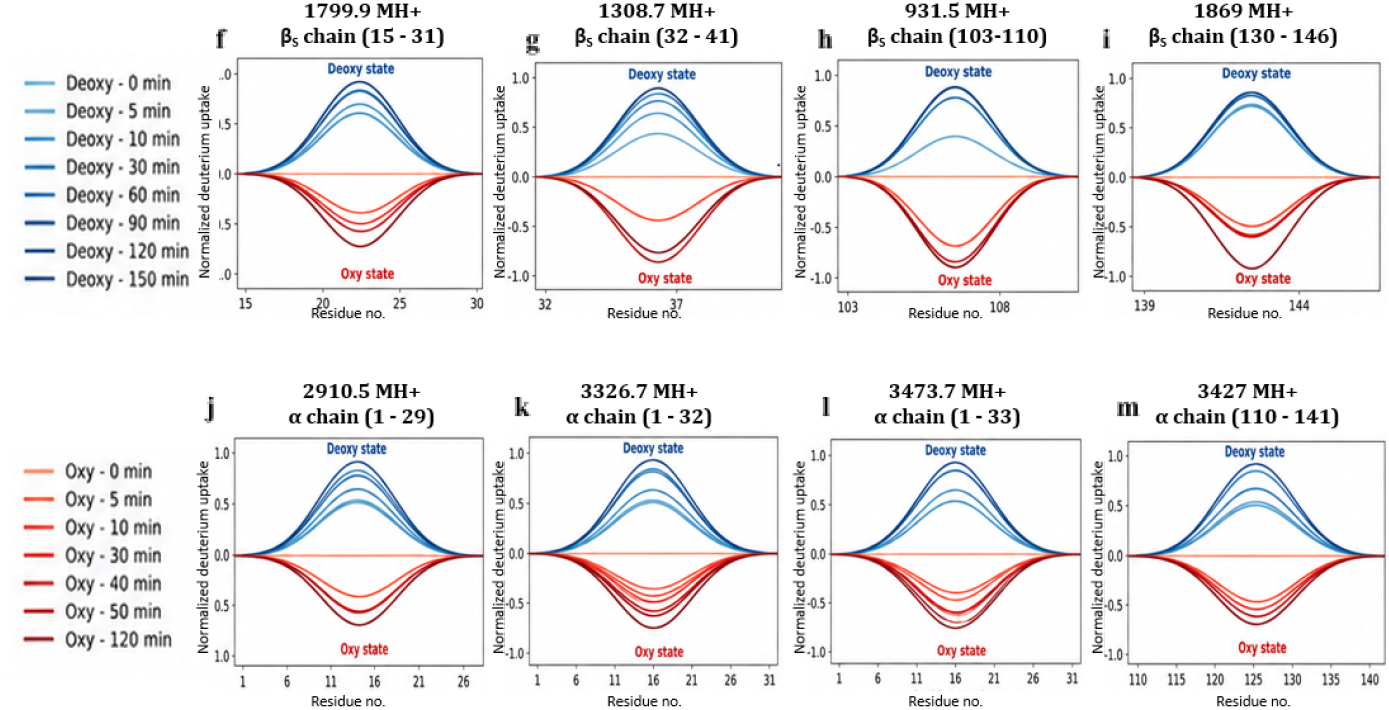
Increase in the isotope-averaged centroid mass across different peptides in oxy and deoxy states: (a-e) Mass spectrum of two different representative peptides (one from α and another from β_S_ chain). In the isotope exchange kinetics, the increasing isotope-averaged centroid mass of the envelope with the increasing time indicates the increased deuterium uptake with time. Panels A and B shows the mass spectra of the deoxy states of both peptides, while Panels C and D are the mass spectra for the oxy state. (f-m) Butterfly plots of the relative deuterium uptake for eight different peptides (four from the α chain and four from the β_S_ chain) for different time points in deoxy (blue) and oxy (red) state.

**Figure 5:**
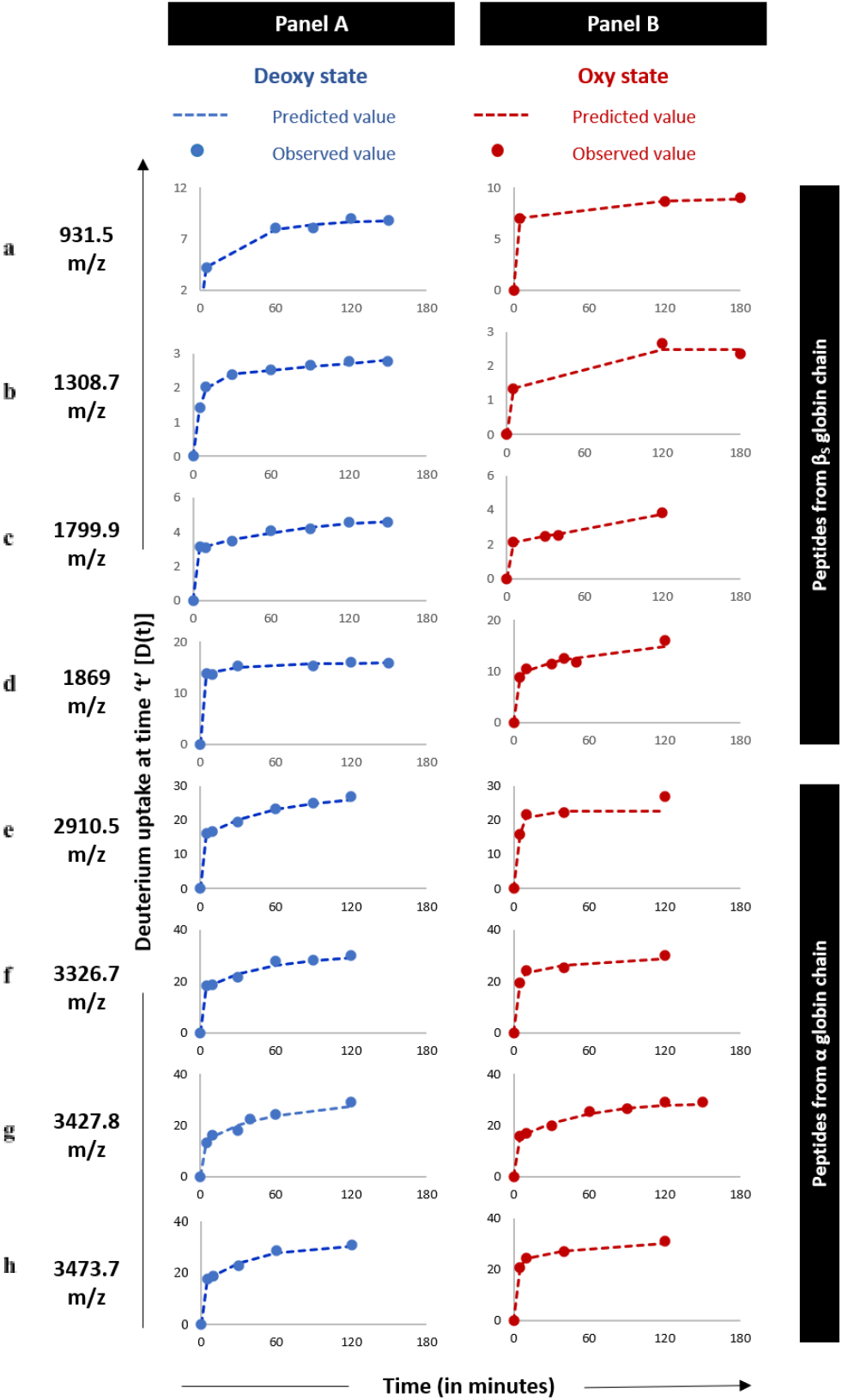
The observed and predicted deuterium uptake plots after fitting the values according to equation 3 in order to minimize the SSR by varying the populations and their respective rate constants for different groups of exchangeable amide hydrogens. (a-h) Panel A shows the uptake plots for the deoxy state, and Panel B shows the uptake plots for the Oxy state.

**Table 1:**
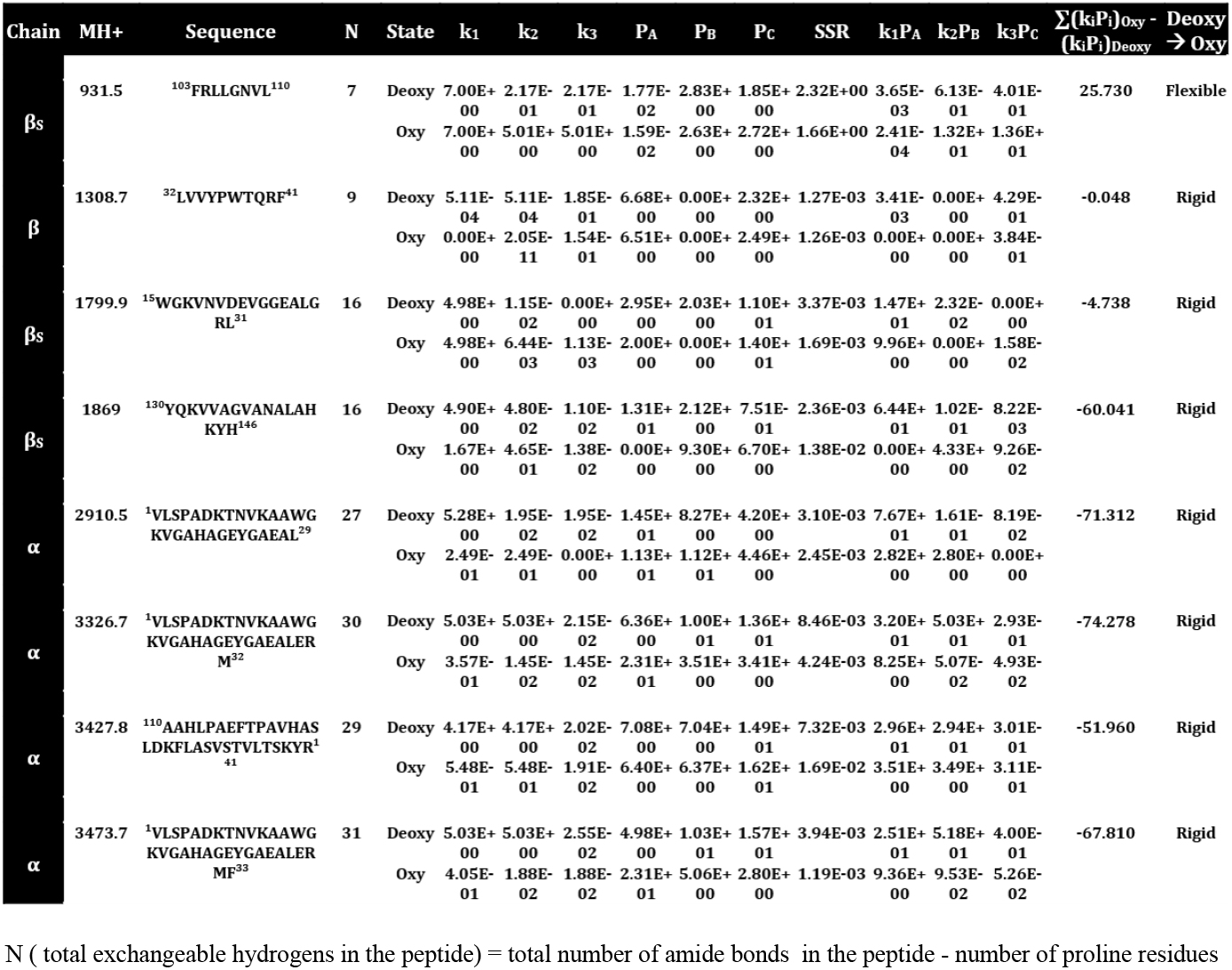
H/DX kinetic parameters of peptic peptides of oxy and deoxy states of sickle hemoglobin.

In the H/DX-MS based study of the experimental sample, four proteolytic peptides from each of the β_S_ and α globin chains were observed, with the following m/z: 931.5 (β_S_ ^103^FRLLGNVL^110^), 1308.7 (β_S_^32^ LVVYPWTQRF^41^), 1799.9 (β_S_^15^ WGKVNVDEVGGEALGRL^31^), 1869 (β_S_^130^ YQKVVAGVANALAHKYH^146^), 2910 (α^1^VLSPADKTNVKAAWGKVGAHAGEYGAEAL^29^), 3326.7 (α^1^VLSPADKTNVKAAWGKVGAHAGEYGAEALERM^32^), 3473.7 (α^1^VLSPADKTNVKAAWGKVGAHAGEYGAEALERMF^33^) and 3427.8 (α^110^AAHLPAEFTPAVHASLDKFLASVSTVLTSKYR^141^).

The peptide with m/z 1308.7 has an identical sequence across the β_S_32-41, γ_G_32-41, and γ_A_32-41 globin chains. Assuming that a peptide sequence, which is shared among different globin chains, might have different local environments, the isotope exchange (H/DX) rates are expected to be different. This should result in a bimodal or multimodal distribution during the exchange kinetics, as the conformations in the respective regions of different tetramers are expected to differ. This, however, was not observed for the peptide with m/z 1308.7, suggesting that it has a similar conformation across the β_S_ and γ globin chains while they are part of different tetrameric species (HbS, HbF, and HbFS). The deoxy to oxy transition in this peptide showed almost no change in flexibility (∑(k_i_P_i_)_Oxy_-(k_i_P_i_)_Deoxy_ **=** -0.048), indicating that the conformation remains protected in this region while transitioning from deoxy to oxy states across all different tetrameric species (HbS, HbF, and HbFS).

The peptide with m/z 931.5 (β_S_103-110) shows a conformational change towards a more flexible state (∑(k_i_P_i_)_Oxy_-(k_i_P_i_)_Deoxy_ = +25.73) in the oxy state compared to the deoxy state. This finding is in agreement with the conformational changes upon deoxy to oxy transitions observed in the HbS (α_2_β_S(2)_) (2), where the overall tetramer becomes more flexible in the oxy state, which does not undergo polymerization.

The peptides with m/z 1799.9 (β_S_15-31) showed a small decrease in the conformational flexibility (∑(k_i_P_i_)_Oxy_-(k_i_P_i_)_Deoxy_ = -4.73), whereas the peptide with m/z 1869 (β_S_130-146) showed a large decrease in its conformational flexibility (∑(k_i_P_i_)_Oxy_-(k_i_P_i_)_Deoxy_ = -60.04) upon deoxy to oxy transition. These peptides consist of various residues that are involved in the polymerization of deoxy HbS. The residues β_S_Gly16, β_S_Lys17, β_S_Val18 and β_S_Glu22 in the peptide with m/z 1799.9 are involved in the axial contacts within the paired strands of deoxy HbS fiber. The residue β_S_Lys144 in the peptide with m/z 1869 has been reported to undergo ion pair interactions with αAsp85, an important contact between two parallel strands of deoxy HbS (36,37). According to our previously reported data, the oxy state exhibits an increased flexibility in these regions, thereby inhibiting HbS polymerization (36). The H/DX data of this study (Table 1) suggest an increase in flexibility of these peptides in the deoxy state compared to the oxy state of HbFS, asymmetric tetramer. This may result in lack of appropriate associations among deoxy HbFS tetramers that favours polymerization and subsequent sickling of RBCs.

All four peptides of the α globin chain with m/z 2910.5, 3326.7, 3473.7 and 3427.8 adopted a more rigid conformation in the oxy state as compared to the deoxy state, as evidenced by the large negative differential rate of isotope exchange (∑(k_i_P_i_)_Oxy_-(k_i_P_i_)_Deoxy_ = -71.3, -74.27, -67.8, and -51.9, respectively). The N-terminal α globin peptides with m/z 2910.5 (α1-29), 3326.7 (α1-32) and 3473.7 (α1-33) showed a more overall rigid conformation in the oxy state as compared to the C-terminal (α110-141). During polymerization of deoxy HbS, N-terminal residue αLys16 interacts axially with αPro114 and αAla115 residues belonging to the a C-terminal peptide 3472.8 m/z (α110-141) of a second tetramer (36, 37). Additionally, the C-terminal residues αPro114 and αAla115 of two different tetramers also interact axially in the HbS fibers (37). Thus, a large increase in the conformational rigidity upon deoxy to oxy transition across this region of HbS might indicate that the necessary interactions that are required for polymerization of the HbS in the deoxy state might not form adequately in the HbFS asymmetric tetramers.

The molecular insights of the inhibition of polymerization contributed by asymmetric hemoglobin referred in this study could have been explored in more detail if it is possible to isolate HbFS devoid of HbS and HbF. Since HbFS is always in dynamic equilibrium with HbF and HbS in a clinical sample obtained from a patient with SCD. Therefore, it is not feasible to isolate the asymmetric tetramer HbFS from the HbS tetramer using standard analytical platforms. However, with the recent advancements in the resolution that can be attained using cryo-electron microscopy (Cryo-EM), it might be possible to resolve the structural intricacies of the HbFS tetramer even in the presence of HbF and HbS. This might provide us with additional insights into the structural aspects to build a more detailed mechanistic model of inhibition in the polymerization of deoxy HbS in the presence of the asymmetric hemoglobin, HbFS.

## Conclusion

The overall conformational state of the HbFS asymmetric tetramer becomes more flexible in the deoxy state compared to the oxy state, thereby lacking the formation of important interactions that are required to stabilize HbS polymers. This might therefore be the molecular basis of the observed inhibition in polymerization of HbS in the presence of the elevated HbF, which eventually reduced the sickling of RBCs.

## Acknowledgments

We acknowledge the funding support from the Indian Council of Medical Research (ICMR), Government of India (Project no. 56/01/2022-Hae/BMS) to carry out this project. We also acknowledge Department of Science and Technology (DST), India, for providing funding for the mass spectrometric facility sanctioned under the project: SR/NM/NS-1068/2015.

## References

1. Pauling L, Itano HA. Sickle cell anemia a molecular disease. Science. 1949 Nov 25;110(2865):543–8. doi:10.1126/science.110.2865.543 PubMed PMID: 15395398.

2. Das R, Mitra A, Bhat V, Mandal AK. Application of isotope exchange based mass spectrometry to understand the mechanism of inhibition of sickle hemoglobin polymerization upon oxygenation. J Struct Biol. 2017 Jul;199(1):76–83. doi:10.1016/j.jsb.2017.04.009 PubMed PMID: 28465180.

3. Eaton WA. Hemoglobin S polymerization and sickle cell disease: A retrospective on the occasion of the 70th anniversary of Pauling’s Science paper. Am J Hematol. 2020 Feb;95(2):205–11. doi:10.1002/ajh.25687 PubMed PMID: 31763707; PubMed Central PMCID: PMC7003899.

4. Wilson SM, Makinen MW. Electron microscope study of the kinetics of the fiber-to-crystal transition of sickle cell hemoglobin. Proc Natl Acad Sci U S A. 1980 Feb;77(2):944–8. doi:10.1073/pnas.77.2.944 PubMed PMID: 6928690; PubMed Central PMCID: PMC348399.

5. Waterman MR, Cottam GL. Molecular Aspects of Sickle Cell Disease. Angew Chem Int Ed Engl. 1976;15(12):749–57. doi:10.1002/anie.197607491

6. Hoffman R, Benz EJ, Silberstein LE, Heslop HE, Weitz JI, Anastasi J, et al. Hematology: Basic Principles and Practice [Internet]. Elsevier Inc.; 2017 [cited 2026 Jun 3]. Available from: https://www.scopus.com/pages/publications/85055776274 doi:10.1016/C2013-0-23355-9

7. Hofrichter J, Ross PD, Eaton WA. Kinetics and Mechanism of Deoxyhemoglobin S Gelation: A New Approach to Understanding Sickle Cell Disease* [Internet]. 1974 [cited 2026 Jun 3]. Available from: https://www.pnas.org/doi/10.1073/pnas.71.12.4864 doi:10.1073/pnas.71.12.4864

8. Mandal AK, Mitra A, Das R. Sickle Cell Hemoglobin. Subcell Biochem. 2020;94:297–322. doi:10.1007/978-3-030-41769-7_12 PubMed PMID: 32189305.

9. Cohen AE, Mahadevan L. Kinks, rings, and rackets in filamentous structures. Proc Natl Acad Sci U S A. 2003 Oct 14;100(21):12141–6. doi:10.1073/pnas.1534600100 PubMed PMID: 14530401; PubMed Central PMCID: PMC218726.

10. Ferrone FA, Hofrichter J, Eaton WA. Kinetics of sickle hemoglobin polymerization: II. A double nucleation mechanism. J Mol Biol. 1985 Jun 25;183(4):611–31. doi:10.1016/0022-2836(85)90175-5

11. Lu L, Li Z, Li H, Li X, Vekilov PG, Karniadakis GE. Quantitative prediction of erythrocyte sickling for the development of advanced sickle cell therapies. Sci Adv. 2019 Aug 21;5(8):eaax3905. doi:10.1126/sciadv.aax3905 PubMed PMID: 31457104; PubMed Central PMCID: PMC6703859.

12. Elendu C, Amaechi DC, Alakwe-Ojimba CE, Elendu TC, Elendu RC, Ayabazu CP, et al. Understanding Sickle cell disease: Causes, symptoms, and treatment options. Medicine (Baltimore). 2023 Sep 22;102(38):e35237. doi:10.1097/MD.0000000000035237 PubMed PMID: 37746969; PubMed Central PMCID: PMC10519513.

13. Naik RP, Whitley KS, Hassell KL, Umeh NI. Clinical Outcomes Associated With Sickle Cell Trait: A Systematic Review: Annals of Internal Medicine: Vol 169, No 9 [Internet]. 2018 [cited 2026 Jun 3]. Available from: https://www.acpjournals.org/doi/10.7326/M18-1161

14. Sachdev V, Rosing DR, Thein SL. Cardiovascular complications of sickle cell disease. Trends Cardiovasc Med. 2021 Apr;31(3):187–93. doi:10.1016/j.tcm.2020.02.002 PubMed PMID: 32139143; PubMed Central PMCID: PMC7417280.

15. Rees DC, Brousse VAM, Brewin JN. Determinants of severity in sickle cell disease. Blood Rev. 2022 Nov 1;56:100983. doi:10.1016/j.blre.2022.100983

16. Tewari S, Brousse V, Piel FB, Menzel S, Rees DC. Environmental determinants of severity in sickle cell disease | Haematologica [Internet]. 2015 [cited 2026 Jun 3]. Available from: https://haematologica.org/article/view/7485

17. Akinsheye I, Alsultan A, Solovieff N, Ngo D, Baldwin CT, Sebastiani P, et al. Fetal hemoglobin in sickle cell anemia. Blood. 2011 Jul 7;118(1):19–27. doi:10.1182/blood-2011-03-325258 PubMed PMID: 21490337; PubMed Central PMCID: PMC3139383.

18. Akinsheye I, Solovieff N, Ngo D, Malek A, Sebastiani P, Steinberg MH, et al. Fetal hemoglobin in sickle cell anemia: molecular characterization of the unusually high fetal hemoglobin phenotype in African Americans. Am J Hematol. 2012 Feb;87(2):217–9. doi:10.1002/ajh.22221 PubMed PMID: 22139998; PubMed Central PMCID: PMC3302931.

19. Bookchin RM, Nagel RL, Balazs T. Role of hybrid tetramer formation in gelation of haemoglobin S. Nature. 1975 Aug;256(5519):667–8. doi:10.1038/256667a0

20. Mtatiro SN, Makani J, Mmbando B, Thein SL, Menzel S, Cox SE. Genetic variants at HbF-modifier loci moderate anemia and leukocytosis in sickle cell disease in Tanzania. Am J Hematol. 2015 Jan;90(1):E1–4. doi:10.1002/ajh.23859 PubMed PMID: 25263325; PubMed Central PMCID: PMC4737118.

21. Steinberg MH. Fetal Hemoglobin in Sickle Hemoglobinopathies: High HbF Genotypes and Phenotypes. J Clin Med. 2020 Nov 23;9(11):3782. doi:10.3390/jcm9113782 PubMed PMID: 33238542; PubMed Central PMCID: PMC7700170.

22. Kaufman DP, Khattar J, Lappin SL. Physiology, Fetal Hemoglobin. In: StatPearls [Internet]. Treasure Island (FL): StatPearls Publishing; 2026 [cited 2026 Jun 3]. Available from: http://www.ncbi.nlm.nih.gov/books/NBK500011/ PubMed PMID: 29763187.

23. Ngo DA, Steinberg MH. Genomic approaches to identifying targets for treating β hemoglobinopathies. BMC Med Genomics. 2015 Jul 29;8:44. doi:10.1186/s12920-015-0120-2 PubMed PMID: 26215470; PubMed Central PMCID: PMC4517356.

24. Platt OS, Brambilla DJ, Rosse WF, Milner PF, Castro O, Steinberg MH, et al. Mortality in sickle cell disease. Life expectancy and risk factors for early death. N Engl J Med. 1994 Jun 9;330(23):1639–44. doi:10.1056/NEJM199406093302303 PubMed PMID: 7993409.

25. Eaton WA. Impact of hemoglobin biophysical studies on molecular pathogenesis and drug therapy for sickle cell disease. Mol Aspects Med. 2022 Apr;84:100971. doi:10.1016/j.mam.2021.100971 PubMed PMID: 34274158; PubMed Central PMCID: PMC8758802.

26. Eaton WA, Bunn HF. Treating sickle cell disease by targeting HbS polymerization. Blood. 2017 May 18;129(20):2719–26. doi:10.1182/blood-2017-02-765891 PubMed PMID: 28385699; PubMed Central PMCID: PMC5437829.

27. Kang HA, Barner JC, Lawson KA, Rascati K, Mignacca RC. Impact of adherence to hydroxyurea on health outcomes among patients with sickle cell disease. Am J Hematol. 2023 Jan;98(1):90–101. doi:10.1002/ajh.26765 PubMed PMID: 36251408.

28. Paikari A, Sheehan VA. Fetal haemoglobin induction in sickle cell disease. Br J Haematol. 2018 Jan;180(2):189–200. doi:10.1111/bjh.15021 PubMed PMID: 29143315; PubMed Central PMCID: PMC5898646.

29. Steinberg MH, Lu ZH, Barton FB, Terrin ML, Charache S, Dover GJ. Fetal hemoglobin in sickle cell anemia: determinants of response to hydroxyurea. Multicenter Study of Hydroxyurea. Blood. 1997 Feb 1;89(3):1078–88. PubMed PMID: 9028341.

30. Ofori-Acquah SF, Green BN, Davies SC, Nicolaides KH, Serjeant GR, Layton DM. Mass spectral analysis of asymmetric hemoglobin hybrids: demonstration of Hb FS (alpha2gammabetaS) in sickle cell disease. Anal Biochem. 2001 Nov 1;298(1):76–82. doi:10.1006/abio.2001.5358 PubMed PMID: 11673898.

31. Rotter M, Aprelev A, Adachi K, Ferrone FA. Molecular Crowding Limits the Role of Fetal Hemoglobin in Therapy for Sickle Cell Disease. J Mol Biol. 2005 Apr 15;347(5):1015–23. doi:10.1016/j.jmb.2005.02.006

32. Yoon YZ, Hong H, Brown A, Kim DC, Kang DJ, Lew VL, et al. Flickering Analysis of Erythrocyte Mechanical Properties: Dependence on Oxygenation Level, Cell Shape, and Hydration Level. Biophys J. 2009 Sep 16;97(6):1606–15. doi:10.1016/j.bpj.2009.06.028 PubMed PMID: 19751665; PubMed Central PMCID: PMC2741588.

33. Ruotolo BT, Benesch JLP, Sandercock AM, Hyung SJ, Robinson CV. Ion mobility-mass spectrometry analysis of large protein complexes. Nat Protoc. 2008;3(7):1139–52. doi:10.1038/nprot.2008.78 PubMed PMID: 18600219.

34. Tuttle LM, James EI, Georgescauld F, Wales TE, Weis DD, Engen JR, et al. Rigorous Analysis of Multimodal HDX-MS Spectra. J Am Soc Mass Spectrom. 2025 Feb 5;36(2):416–23. doi:10.1021/jasms.4c00471 PubMed PMID: 39837577; PubMed Central PMCID: PMC12034455.

35. Srinivasu BY, Mandal A. Application of Hydrogen / Deuterium Exchange Mass Spectrometry in Structural Biology and Molecular Medicine. In. 2017 [cited 2026 Jun 3]. Available from: https://www.semanticscholar.org/paper/Application-of-Hydrogen-Deuterium-Exchange-Mass-in-Srinivasu-Mandal/37e02677adf5d531cb4793d70d60211720b401bd

36. Das, R., Mitra, A., Bhat, V., & Mandal, A. K. (2017). Application of isotope exchange based mass spectrometry to understand the mechanism of inhibition of sickle hemoglobin polymerization upon oxygenation. Journal of Structural Biology, 199(1), 76–83. 10.1016/j.jsb.2017.04.009

37. Harrington, D. J., Adachi, K., & Royer, W. E. (1997). The high resolution crystal structure of deoxyhemoglobin S1. Journal of Molecular Biology, 272(3), 398–407. 10.1006/jmbi.1997.1253

