## Supplementary figures for "Structural insights into the inhibition of sickle hemoglobin polymerization by asymmetric hemoglobin tetramer HbFS (α_2_γβ_S_)"

a

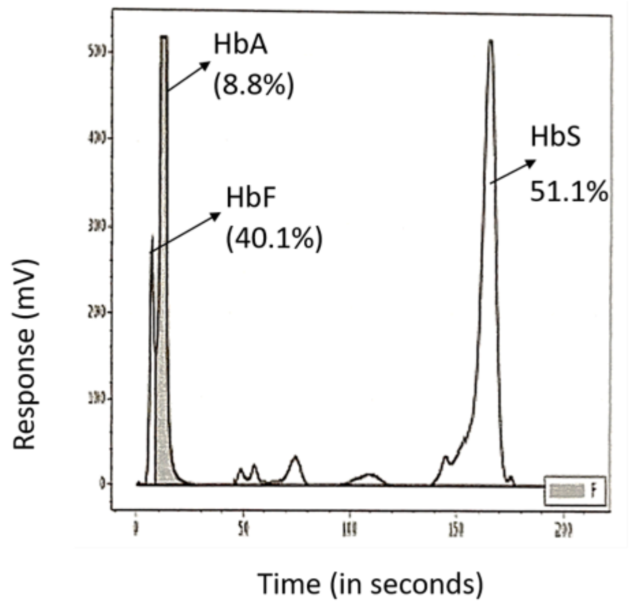


b

| TEST | RESULT | UNIT | Normal range (*Adult) |
|---|---|---|---|
| Haemoglobin A | **8.8** | % | 94.3 - 98.5 % |
| Haemoglobin A2 | 0.0 | % | 1.5 - 3.7 % |
| Haemoglobin F | **40.1** | % | 0.0 - 2.0 % |
| Haemoglobin S | 51.1 | % | 0.0 - 0.0 % |
| Haemoglobin D | 0.0 | % | 0.0 - 0.0 % |
| Haemoglobin C | 0.0 | % | 0.0 - 0.0 % |
| Unknown Peak | 0.0 | % | 0.0 - 2.0 % |

**Figure 1:** HPLC analysis of hemoglobin profile of the patient (a), at the diagnostic settings of AIIMS Kalyani, revealed the presence of HbS, HbF, and HbA at 51.1, 40.1, and 8.8%, respectively (b).

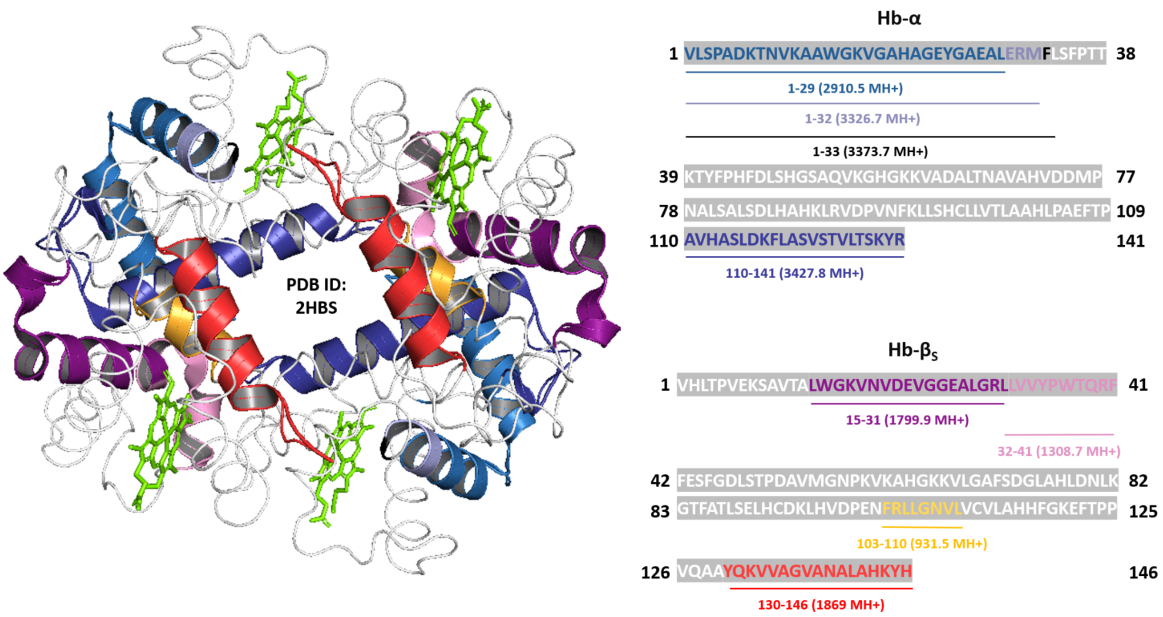


**Figure 2:** The peptides identified across the α and $\beta_S$ chains in the MALDI spectrum, mapped onto the HbS tetramer crystal structure (PDB Id: 2HBS).

a

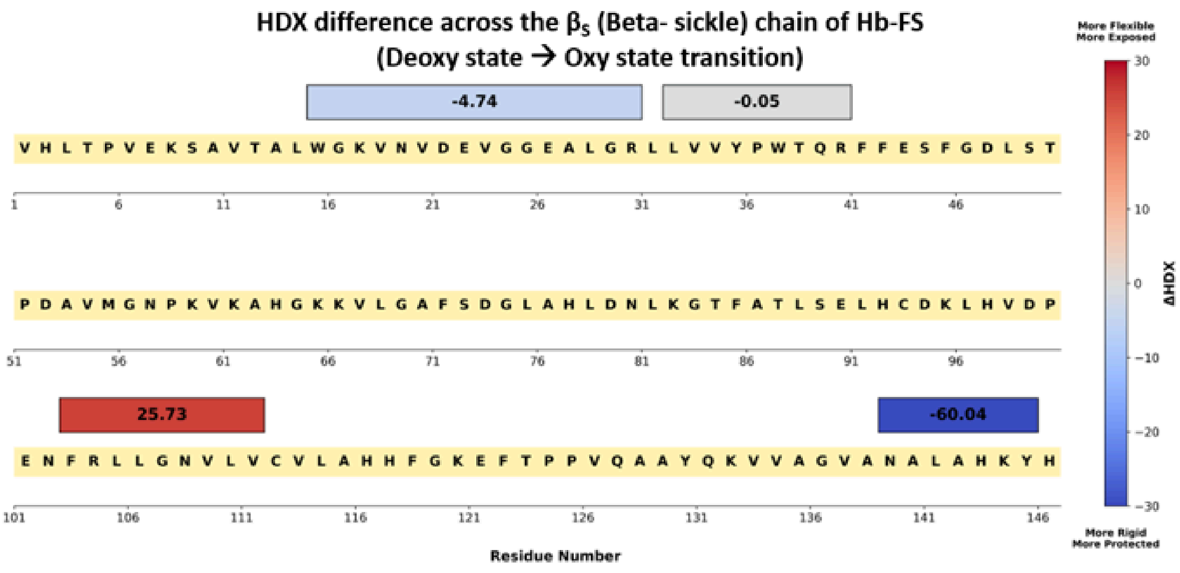


b

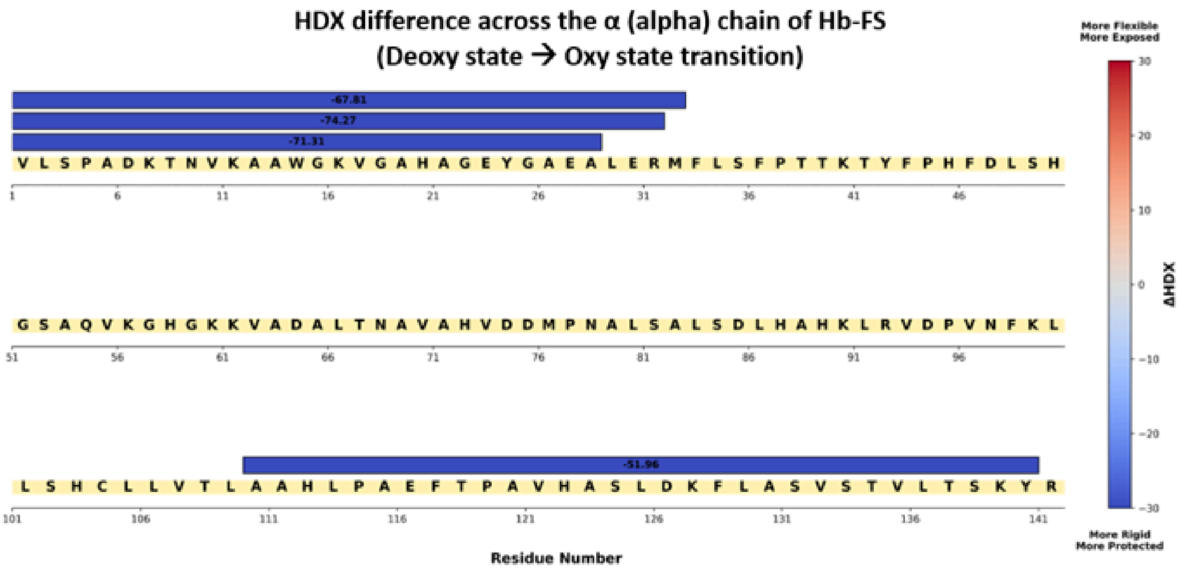


**Figure 3:** The H/DX-MS difference map across the alpha chain (a) and the beta chain (b) of the HbFS sample shows that the deoxy to oxy state transition of the hemoglobin does results in overall rigidity of the both α and $\beta_S$ chains of the tetramer.
